# Beyond High-Amplitude Events: Amplitude-Resolved Spatial and Temporal ICA Representations in fMRI

**DOI:** 10.64898/2026.10.07.757468

**Authors:** Duc Vu, Vince Calhoun

## Abstract

Functional MRI analyses commonly reduce high-dimensional brain activity using predefined spatial summaries, such as region-of-interest (ROI) averages, or temporally selective representations, such as co-activation patterns (CAPs) and high-amplitude events. These approaches provide useful summaries but do not explicitly characterize information distribution across the continuous magnitude of spatial and temporal network expression. We introduce an amplitude-resolved framework for systematically interrogating this continuum using independent component analysis (ICA). Resting-state fMRI data in the FBIRN dataset were decomposed using spatially-constrained ICA with NeuroMark. For the temporal representation, time points were stratified according to global ICA amplitude and frequency bands, with functional network connectivity (FNC) estimated for each amplitude range. For the spatial representation, we introduce an approach that stratifies voxels according to ICA loading magnitude and generates amplitude-specific spatial similarity matrices. Linear support vector machine (SVM) classification of schizophrenia (SZ) and healthy control (HC) participants was used to probe the information retained by each representation. Discriminative information varied non-monotonically across amplitude: intermediate ranges frequently matched or exceeded the highest amplitudes, while combinations of amplitude ranges further improved classification. These findings demonstrate that informative fMRI structure is distributed across the ICA expression continuum and motivate representations that complement fixed ROI summaries and high-amplitude approaches.

## 1. INTRODUCTION

Functional MRI produces high-dimensional signals that vary continuously across both space and time, yet many analyses require substantial summarization before statistical or machine-learning analysis. Spatially, a common strategy is to average signals within predefined regions of interest (ROIs) or parcels[1], producing robust and interpretable regional time courses but reducing within-region spatial variation. Temporally, approaches such as co-activation patterns (CAPs), event-based analyses, and related thresholding methods focus on selected frames or configurations, often emphasizing periods of strong or high-amplitude activity[2]. These approaches have provided important insights into functional brain organization, but they address particular spatial or temporal summaries rather than explicitly asking how information is distributed across the continuous magnitude of network expression.

A complementary question is therefore whether the strongest fMRI signals are necessarily the most informative. High-amplitude events may capture particularly salient network configurations, but biologically or clinically relevant information may also be expressed at intermediate or relatively weak levels of activity[3]. Similarly, spatial information need not be confined to the most strongly expressed voxels within a functional network. Characterizing the full amplitude continuum could therefore reveal structure that is not apparent when signals are reduced to fixed regional summaries or restricted to high-amplitude events.

To investigate the full amplitude range of fMRI signals, we apply spatially constrained ICA using NeuroMark templates [4] on FBIRN dataset [5] and threshold the output into amplitude percentiles. Since ICA outputs voxel-level signals, we can examine a broader spectrum than CAPs approaches or fixed ROIs. Along with conventional time course filtering and our amplitude percentile thresholding, we introduce a new method of obtaining component map percentiles, inspired by maximum intensity projection [6]. With thresholded time courses and component maps, we construct FNCs and similarity matrices as features of an SVM for schizophrenia classification. This framework determines whether focusing primarily on the largest-amplitude events sufficient for schizophrenia identification, or if relevant information spread throughout the continuum of temporal and spatial amplitudes.

## 2. MATERIALS AND DATA PRE-PROCESSING

We used fMRI data from fBIRN project [7] for eye-closed resting-state fMRI scans from 311 subjects in total, with 151 schizoprenia subjects (SZ) and 160 age and gender-matched healthy controls (HC). Informed consent was obtained from the participants. The fMRI data was collected using 3T MRI machines. The imaging parameters were as follows: 64 *×*64 acquisition matrix, repetition time (TR) of 2 s, echo time (TE) of 30 ms, flip angle (FA) of 77°, voxel size of 3.4 *×*3.4*×* 4 mm, slice gap of 1 mm, and 162 acquired volumes.

Data pre-processing included brain extraction, slicetiming, and motion correction. The data was then registered to MNI space, resampled to 3 mm^3^ isotropic voxels and spatially smoothed using a Gaussian kernel with 6mm full-width at half maximum. For each subject’s fMRI data, we applied spatially constrained ICA using the GIFT toolbox [8] with NeuroMark fMRI version 1.0 template [4], resulting in 53 reproducible independent component networks (ICNs) that is partitioned into seven subcategories: subcortical (SC), auditory (AUD), visual (VIS), sensorimotor (SM), cognitive control (CC), default mode network (DMN) and cerebellar (CB). We obtain the component maps **S** *∈ R*^*C×V*^ (Z-scored across all voxels within the brain) and time course matrix **A** *∈ R*^*T ×C*^ (Z-scored across time). C is the number of components, V is the number of voxels within the brain, and T is the number of time points.

## 3. METHODOLOGY

### 3.1. Time Course Percentile and Frequency Thresholding

To isolate signals within selected frequency ranges, we band-pass filtered each time course using a Butterworth filter. We examined (0.01− 0.08, Hz), the conventional amplitude of low frequency fluctuation (ALFF) range; (0.08− 0.15, Hz), a higher BOLD range commonly retained in ICA; (0.15 −0.249, Hz), the upper BOLD-sensitive range; and (0.01 −0.249, Hz), the full bandwidth examined (Fig.1**(d)**) [9, 10].

**Fig. 1.**
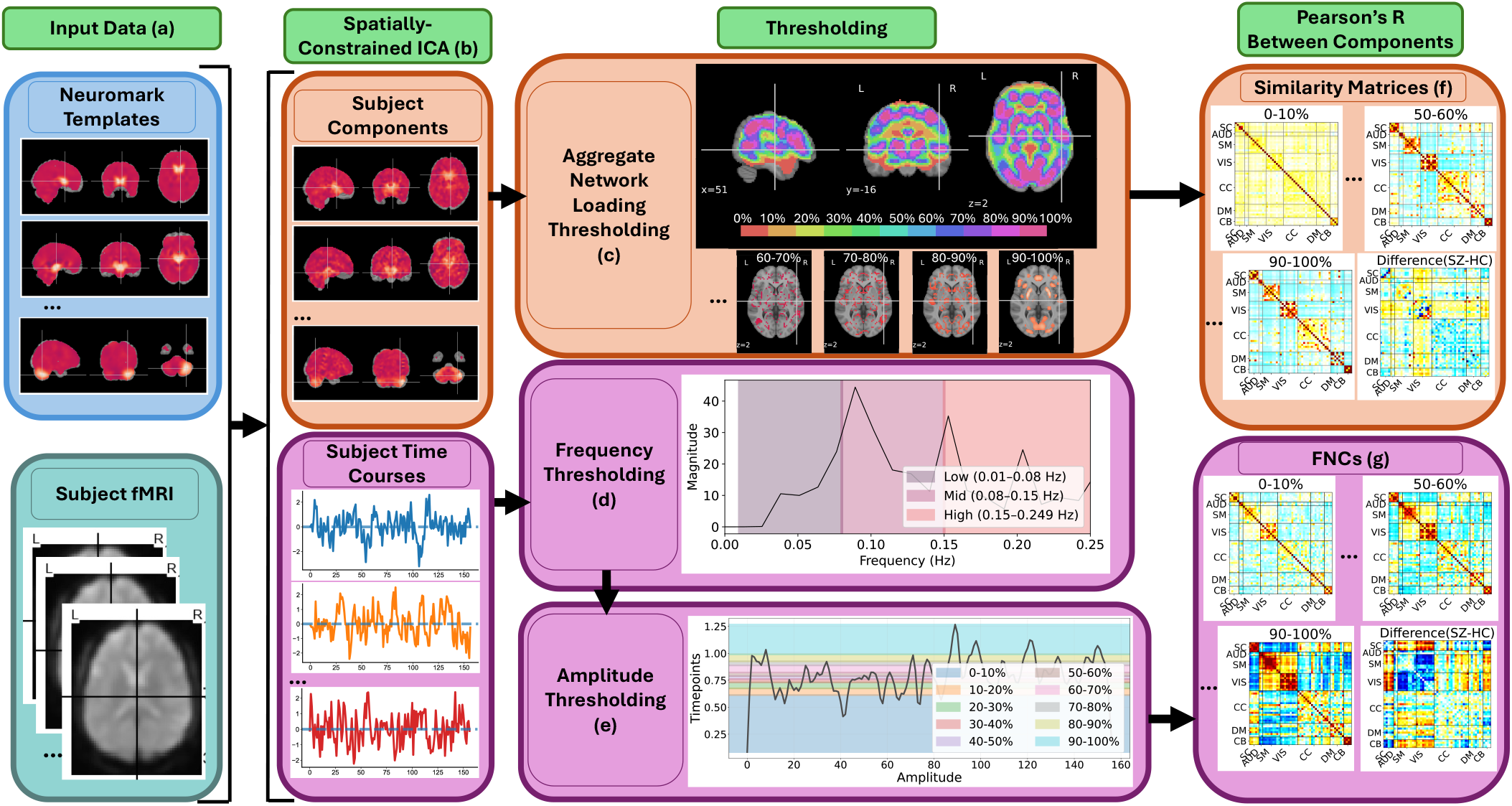
We process the fMRI data with spatially-constrained ICA using Neuromark templates. **(a)**. After obtaining the time course and component maps **(b)**, we threshold each subject’s ICA output with conventional time course frequency and amplitude thresholding **(d)(e)** and Aggregate Network Loading thresholding **(c)** over all components of that subject. From the thresholded signals, we obtain the corresponding thresholded similarity matrix **(f)** and FNC **(g)** as features for discerning schizophrenia. The difference in FNCs and similarity matrices between schizophrenia and healthy control group are also shown in **(f)(g)**.

After filtering, we threshold the time course matrix **A** for each subject. The absolute value of the timecourse for each component was then averaged across component: 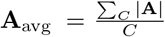. **A**_avg_ can be seen as the average amplitude of components across time. We threshold by taking percentiles with 10% bins (Fig.1 **(e)**) to get the equivalent time points *T*_%_ of each percentiles as masks. We obtain the value of each component’s time course during those *T*_%_ time points and get syn-chronized component time courses 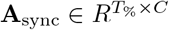. We use **A**_sync_ to construct the FNC of the subject **F** *∈ R*^*C×C*^ (Fig. 1 **(g)**). The figure shows that emphasized inter-network correlations are different for each percentile.

### 3.2. Component Map Aggregate Network Loading

For component maps, we threshold using a new method we call Aggregate Network Loading: for each subject’s component map **S**, we obtain the maximum of each voxel within the brain across components to obtain the maximum intensity projection **V** *∈ R*^*V*^ (Fig.1**(c)**). This shows where the ICA component map signal is strongest across all components of a subject. We then threshold the signal to obtain voxels as binary masks **V**_**mask**_ *∈ R*^*V*^ for each percentiles similar to Fig.1**(c)** masks. With each component C of the subject **S**_**C**_, we use the masks to compute the respective percentile’s component signal **S**_**C**%_ *∈ R*^*V*^. We construct the spatial similarity matrices **F** ∈ *R*^*C×C*^ using Pearson correlation between each pair of component maps **S**_**C**%_ of a subject. Previous work [11][12] has shown that Pearson correlation is often used to match components across ICA methods, and spatial similarity patterns in ICA differ between healthy and schizophrenia subjects.

With these percentile masks, the resulting similarity matrices varies over different percentile ranges for each subject.Fig. 1 **(f)** demonstrates that patterns change with percentiles, with 0 −10% percentile showing mostly noise but we can see that patterns within-network are already showing. More useful patterns show up around the 50 − 60% range and beyond. As expected, components within the same network have higher spatial similarity.

## 4. EXPERIMENT AND RESULTS

### 4.1. Discriminative Time Course Amplitudes

To identify discriminative information in temporal structures, we used the entries of the time course FNCs as features for an SVM model to classify between HC and SZ subjects[13]. The model performed a 10-fold cross validation with the test pool kept completely separate from the training pool to prevent data leakage. The SVM ran 10 times with the reported AUC being the average of those runs (Fig.2**(a)**).

The AUC results vary significantly across different percentiles and frequency bands, with (0.01 − 0.08, Hz) and (0.01 − 0.249, Hz) being most discriminative. For (0.01− 0.08, Hz), the intermediate and low percentiles (30 − 40% and 50 − 60%) performed just as well as high percentiles (80 − 90%) and far exceeds the highest percentile, showing the importance of information residing outside the highest ampli-tudes.

**Fig. 2.**
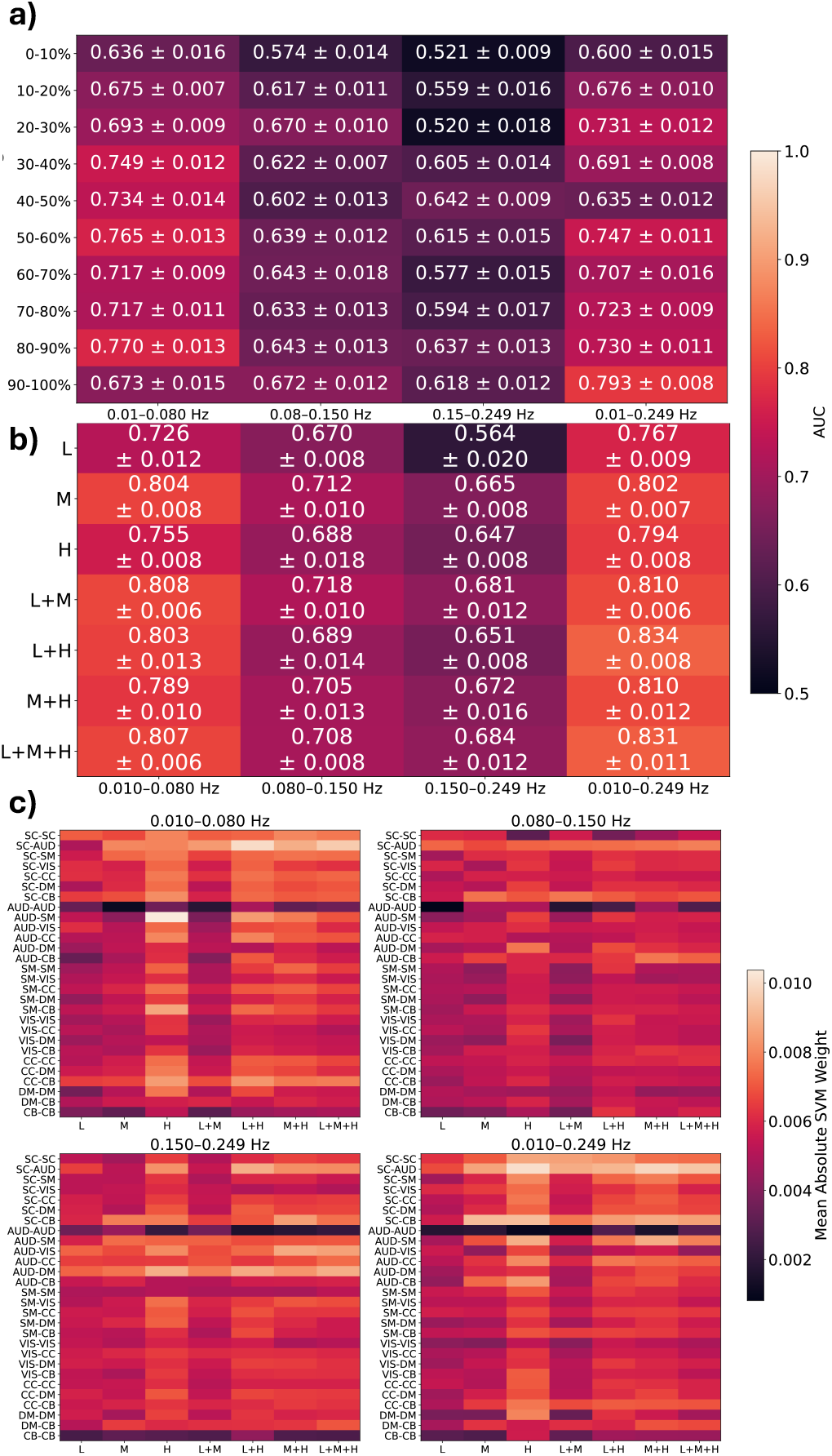
Time course result. **(a)** Result grid of SVM on temporal data using different frequency band-pass filter and percentile thresholding. **(b)** Result grid with percentiles merged into low (L), mid (M), and high (H). **(c)** SVM weight matrix of network pairs for each merging schemes and frequency band.

We determine whether the information in each percentile is complementary by merging the time points selected in different percentiles to construct FNCs and use it as features. We chose our merging scheme as combinations of low (L), mid (M), and high (H) bands: low bands are merges of 0 −30%, mid bands are merges of 30 −70%, and high bands are merges of 70 −100%. The result in Fig.2**(b)** shows that for both discriminative frequency band, the best results are L+M or L+H, with AUC much higher than the highest AUC of Fig.2**(a)**. This demonstrates that information in different percentiles are complementary, and merging low percentiles with mid or high will significantly boost performance, despite low percentiles having the worst AUC in all frequency band.

To quantify the discerning information from each network pair, we computed the absolute values of SVM weights, corresponding to component pairs contribution, and averaged it across runs, subjects, and within networks to obtain network-pair SVM weight in Fig.2**(c)**. The significance of most pairs differ significantly across percentiles and frequency, while some pairs like SC-AUD are consistent in all frequencies. From the frequency band with highest AUC (0.01 −0.249, Hz), we find clear contributions from pairs that are validated by recent research[4][14]: SC-AUD and SC-CB contributed more in mid and high, and AUD-SM, AUD-CB, SC-SC, and SC-AUD contributed most in high percentiles.

To further investigate the consistency of important network pairs, we obtain the difference in FNCs between SZ and HC group (Fig.3**(a)**). The figure shows clear differences in inter-network correlations between SC, AUD, CB, VIS, and SM, agreeing with our results and previous findings of schizophrenia subjects’ deficiency in sensory processing[14].

**Fig. 3.**
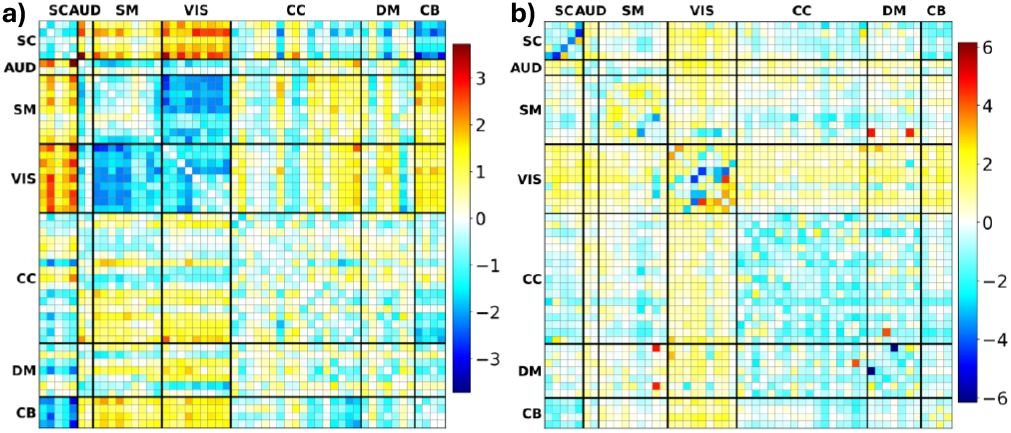
The zscore difference in similarity matrix between schizophrenia and healthy group, averaged over all subjects in each group. **(a)** FNC difference, **(b)** Spatial difference

### 4.2. Discerning Component Map Amplitudes

For component maps, we also used entries of the similarity matrices as features for the SVM in classification. The SVM also performed a 10-fold cross validation for each percentile of spatial data thresholded.

The result grid in Fig.4**(a)** shows a more linear pattern compared to time course, with the highest AUC concentrated around mid-high percentiles (70 −80%) rather than the highest percentile. This indicates that not all discerning information are found in the highest spatial amplitudes.

For complementary information analysis, we employ the same merging scheme as time course analysis. Fig. 4 **(b)** shows AUC results for L, M, and H can be boosted by combining merges, achieving the highest AUC with M+H, higher than Fig. 4 **(a)**. This also suggests supplementary information between spatial amplitude percentiles. However, the AUC for L, M, and H is lower than the individual highest of each merge in **(a)**, implying focused information in smaller bands.

**Fig. 4.**
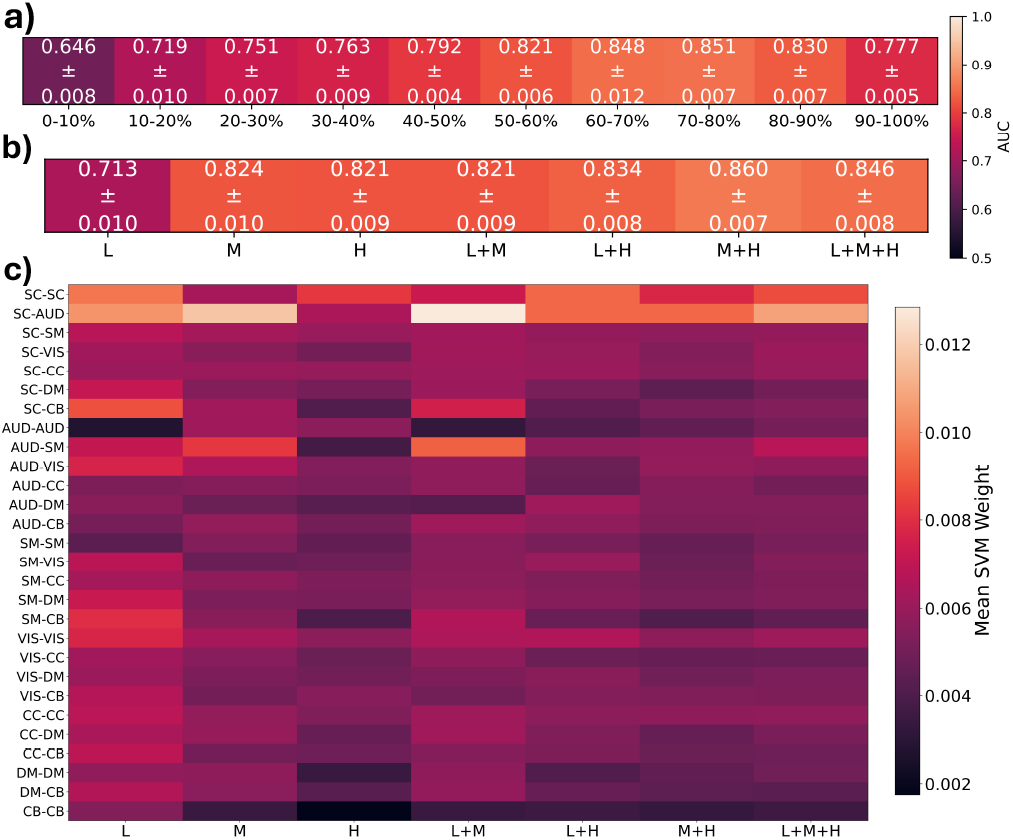
Component map result. **(a)** Result grid of running SVM on percentile spatial similarity matrices. **(b)** Result grid with each percentile merging scheme. **(c)** SVM weight matrix of the merged percentiles.

We construct the SVM weight plot in Fig.4**(c)** similarly to time course using the similarity matrices’ entries. SC-SC and SC-AUD are now prominent network pairs used for differentiating SZ from HC across all percentiles, which were not as obvious in time course data. The difference in spatial similarity matrix between SZ and HC group (Fig. 3 **(b)**) also shows within-network differences of SC (particularly thalamus and caudate) and VIS being discerning, aligning with results in previous work [14].

## 5. CONCLUSION

This paper introduces an amplitude-resolved framework that identifies where discerning information reside throughout time course and component map amplitudes. We introduced a new method to threshold component maps called Aggregate Network Loading and thresholded component maps and time courses to generate similarity matrices and FNCs for SVM classification. The result indicates discriminative information in not just high but sometimes mid and low percentiles, suggesting information at different levels of amplitudes. Merging the percentiles further improves the performance, supporting the complementary nature of these amplitude percentiles. Identified valuable networks are verified by previous research, with certain networks like SM and CB contributing more in higher percentiles. The discriminative information obtained from assessing these moderate and low percentiles can complement CAPs or peak-event approaches in future integrations. Current limitations are lack of interpretations and identification of best combination of percentiles.

## Notes

### Competing Interest Statement

The authors have declared no competing interest.

### Funding

- National Institute of Health: NIH R01EB006841

